# Using a simplified Rough Mount Fuji model to disentangle how multi-peaked fitness landscapes can be highly navigable

**DOI:** 10.64898/2026.03.19.712707

**Authors:** Kye E. Hunter, Nora S. Martin

## Abstract

Evolving populations, especially in the strong-selection-weak-mutation limit, can be modelled as adaptive walks on fitness landscapes, moving in fitness-increasing mutational steps until reaching a fitness peak—a local optimum. Simulations of such adaptive walks—on a multi-peaked empirical landscape of the *folA* gene and on landscapes generated by the ‘Rough Mount Fuji’ (RMF) model— have shown that some landscapes are highly navigable, meaning that the highest *x*% of peaks are reached by ≫ *x*% of adaptive walks. This prompts the question of how adaptive walks can be so successful despite the local, ‘myopic’ rules behind each adaptive step. Here, we investigate this question using simulations and mathematical approximations of random adaptive walks on a simplified RMF landscape. The landscape has a low-to-intermediate fitness region, whose size reconciles a low peak density with a high peak number. Despite the high number of peaks, walkers are likely to exit this region without terminating at a peak because the probability of a peak transition at each step is low and a fitness gradient guides walkers to the high-fitness region in few steps. Thus, three features are sufficient to explain why adaptive walks in the simplified RMF landscape are likely to reach a small fraction of top-ranking peaks: a low-to-intermediate fitness region with a high number of peaks, a low peak-transition probability, and which is crossed in few steps. We find that these three features are also present in the empirical *folA* landscape, suggesting that similar principles may apply.

## I. INTRODUCTION

In biological evolution, fitness-increasing mutations are favoured by natural selection, thus increasing their chances of becoming prevalent in future generations. This can be modelled as populations moving on a fitness landscape, a network, where each node stands for a genotype, its elevation for its fitness, and edges connect two genotypes if one can be reached from the other in a single mutation [1, 2]. In this picture, populations are localised on one or more nodes in the landscape, mutations move individuals to neighbouring nodes, and selection tends to concentrate the population on higher-fitness nodes. This picture is further simplified in the strong-selection-weak-mutation regime, where evolving populations tend to be concentrated on one node at a time and can be approximated as a single adaptive walker [1]. This walker moves through mutational steps along fitness-increasing, *accessible* paths [1] until it ends at a local peak, where no single mutations provide further fitness gains. Thus, fitness peaks constitute the possible end points of adaptive walks, prompting questions about their density, their height and their probability of being realised.

Answering these questions requires a dataset defining a fitness landscape, i.e. a dataset mapping all possible genotypes of length *L* to their respective fitness. Here, we focus on the results from an empirical dataset, covering almost 99.7% of all possible genotypes in a nine-nucleotide section of the *folA* gene in *E. coli* under trimethoprim exposure [3]. After extracting the regions with functional genotypes, Papkou et al. [3] found that this sublandscape has 514 fitness peaks. Among these, simulated adaptive walks end at the highest-14% of peaks with a probability of *>* 75% [3] (Fig. 1A). Thus, adaptive walkers are likely to end at a small subset of the highest peaks on the *folA* landscape, even though the walkers choose their steps probabilistically based only on their immediate mutational neighbours and can thus be described as “myopic” [4] or short-sighted. This is not simply equivalent to reaching high-fitness peaks – which could be achieved in a landscape with *only* high-fitness peaks. Rather, low-fitness peaks are present, but under-represented in adaptive walk endpoints.

**Figure 1.**
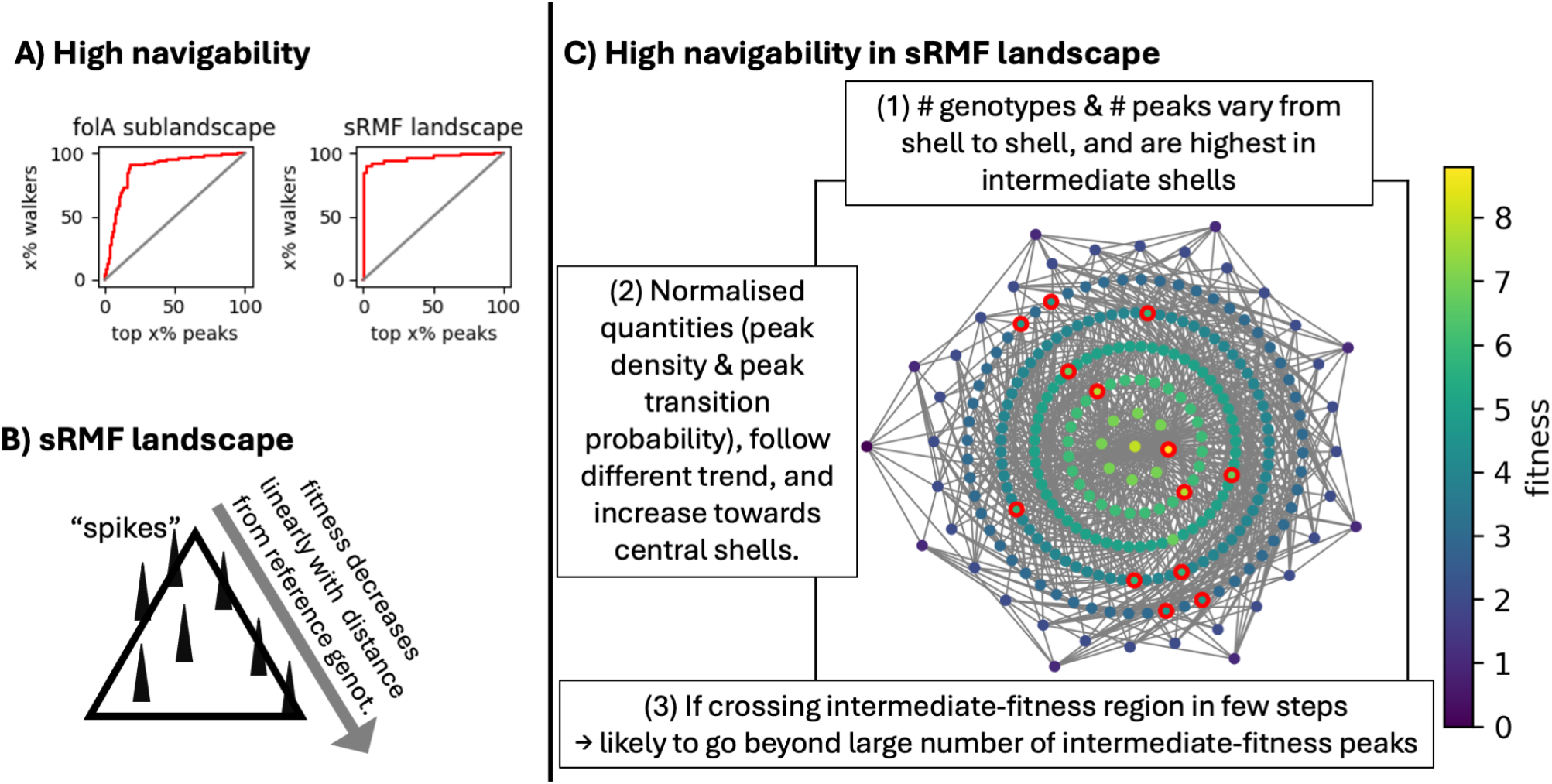
High-navigability phenomenon and our explanation in the sRMF model: (A) Both the empirical *folA* sublandscape and the mathematical sRMF landscape (with *L* = 15, *p* = 0.01, Δ = 1.5) are highly navigable: the highest *x*% of peaks are reached by ≫ *x*% of simulated adaptive walks (red, based on 10^4^ walks). The grey *x* = *y* lines are shown for context. (B) The sRMF model can be visualised as a smooth cone (additive component) with spikes (random component), but this analogy neglects the high dimensionality of genotype space. (C) The sRMF landscape is more accurately represented as a network (here *L* = 8, *p* = 0.05, Δ = 1.8 for clearer visualisation), with genotypes as nodes, genotype fitness in colour and mutations as edges. Genotypes are sorted into shells, depending on their Hamming distance *d* to the reference genotype. Red circles denote peaks, illustrating that a spike is typically a peak unless it is adjacent to a higher-fitness spike. Three features, labelled (1)-(3) in the schematic, are key to our explanation of how the highest *x*% of peaks are reached by ≫ *x*% of adaptive walks. Note that both the *high* peak number and the *low* peak transition probability in the low-to-intermediate fitness region are important: a lower peak number would affect the fitness percentile of the high-fitness peaks, and a higher peak transition probability would affect the probability of reaching these high-fitness peaks.

Explaining this apparent paradox means investigating how a landscape can be structured such that the top *x*% of peaks are reached by ≫*x*% of adaptive walkers – a phenomenon we will refer to as the *high-navigability phenomenon*, thus reserving *navigability* for the *probabilities* of reaching peaks on adaptive walks, in contrast to *accessibility*, which refers to the *existence* of fitness-increasing paths^1^. The high-navigability phenomenon is relevant beyond this specific *folA* landscape since it has been reported in further empirical landscapes, for example the bacterial transcription factor TetR [6] and the SARS-CoV-2 spike RBD [7].

A first step towards understanding this high-navigability phenomenon was made by Li & Zhang [7], who identified a highly navigable landscape that is simpler to analyse: the ‘Rough-Mount-Fuji’ (RMF) model, an abstract mathematical model [8, 9] that reproduces further features of empirical fitness landscapes [9–11]. The RMF model is defined as follows: As a basis, one builds an additive landscape, where each mutation away from a fixed reference genotype has a fixed fitness penalty, and these fitness penalties accumulate additively [12]. To turn this smooth, additive landscape into a *Rough* Mount Fuji, a random value, drawn from a fixed probability distribution, is added to each genotype’s additive fitness [12]. Due to this random component, a genotype’s fitness can surpass all its single-mutant neighbours and thus become a local peak, making the RMF model multi-peaked [9]. Thus, in an RMF landscape, fitness tends to increase with decreasing distance to the reference genotype, but local peaks exist due to random fluctuations (schematic in Fig. 1B).

Several features that may be relevant for the RMF landscapes’ high navigability have been characterised computationally and, in some cases, analytically: First, for RMF models with random components from a Gumbel-type distribution, the density of peaks decreases with increasing distance from the reference genotype [9]. Secondly, RMF landscapes typically have at least one accessible mutational path to the reference genotype [13, 14]. Turning to the set of genotypes with an accessible path to a given peak, i.e. the *basin* of that peak, basin sizes in RMF landscapes tend to increase with peak fitness [7]. However, the *existence* of accessible paths to high-fitness peaks is not enough to guarantee that a high peak is reached since a genotype can have accessible paths to multiple peaks [15]. Thus, information beyond accessible paths is needed, describing the trajectories of adaptive walks. In RMF landscapes, such trajectories can be much longer than in uncorrelated, ‘House-of-Cards’-type landscapes [15], and on RMF models dominated by the additive component trajectories are likely to end on the reference genotype [9, 15]. These results are all consistent with the landscape’s high navigability. However, despite the tractability of the model, we are lacking a simple explanation resolving the apparent paradox of the high-navigability phenomenon—how the RMF landscape’s features allow adaptive walkers to avoid low-fitness peaks.

Here, we address this question with a simplified ver-sion of the RMF model (sRMF model). This simplification further strengthens the conceptual and mathematical tractability of the model while maintaining the high-navigability phenomenon for suitably chosen parameters (Fig. 1A). We use analytic calculations and simulations to gain an overview of the sRMF landscape, identifying the following key features (Fig. 1C): a low-to-intermediate-fitness region, which contains a high number of peaks, yet has a low peak density and low peak transition probability, and is crossed in short paths. Thus, a high fraction of walks reach a fitness exceeding the low-to-intermediate-fitness peaks despite their high number. While the mathematical expressions behind this argument only hold for the sRMF model, we identify similar features in the empirical *folA* landscape. Finally, we connect our results to existing work on basins in highly navigable landscapes.

## II. RESULTS

### A. A simpler version of the RMF model reproduces the high-navigability phenomenon

We will use a simplified RMF (sRMF) model, which is simpler to analyse mathematically and conceptually than the standard RMF model, and, while idealised, highly suitable for understanding the high-navigability phenomenon. In the sRMF model, each genotype is a length-*L* sequence chosen from a binary alphabet {0, 1}. Like standard RMF landscapes [9], the sRMF model sets each genotype’s fitness as the sum of an additive fitness component and a random component, where the additive component depends linearly on the Hamming distance *d* of the genotype to a reference genotype (0, 0, 0, …) as *L* − *d*. The random component is simpler than the continuous random variable in standard RMF models and is chosen from two possible discrete values: either Δ, with probability *p* (a *spike* genotype), or zero otherwise (a *non-spike* genotype). This random component in isolation is equivalent to an existing model termed ‘holey landscapes’ [16], while standard RMF models mirror ‘House-of-Cards’ landscapes [9]. Adding the additive and random components, the fitness *f* (*g*) of a genotype *g* is:

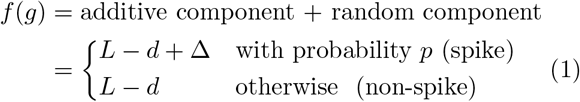

Δ sets the elevation of spike genotypes relative to non-spikes. Multi-peaked landscapes require Δ *>* 1, and for 1 *<* Δ *<* 2 the global maximum will be at most one mutation from the reference genotype. In the following, we choose Δ = 1.5, but other values between 1 *<* Δ *<* 2 would leave fitness ranks unaffected. To simplify calculations, we will focus on the low-*p* limit, where spikes are unlikely to be adjacent to further spikes and thus each spike can be assumed to be a peak. Thus, our simulations take *p* = 0.01, *L* = 15 and Δ = 1.5, except where otherwise stated.

Because fitness could be rescaled arbitrarily in the sRMF model, we focus on *random adaptive walks*, which are not sensitive to absolute fitness values, but only to their rank ordering: every fitness-increasing mutational step is equally likely [17]. These random adaptive walks are sufficient to model the key phenomenon we are interested in: they reach the top-14% peaks on the empirical *folA* landscape with a probability of *>* 70% [3] despite their myopic and stochastic nature.

### B. The shells with the highest number of peaks do not have the highest peak density

Before analysing the peaks reached on adaptive walks, we establish some context: the location and height of all peaks in the sRMF landscape. For this purpose, we will divide our landscape into shells, genotypes with a fixed Hamming distance *d* from the reference genotype and thus the same additive fitness component. The number of genotypes in a shell is given by the number of ways of choosing *d* mutations relative to the reference genotype:

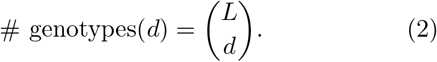

Thus, the medium-fitness shells with *d* ∼*L/*2 contain the highest number of genotypes, see Fig 2A.

**Figure 2.**
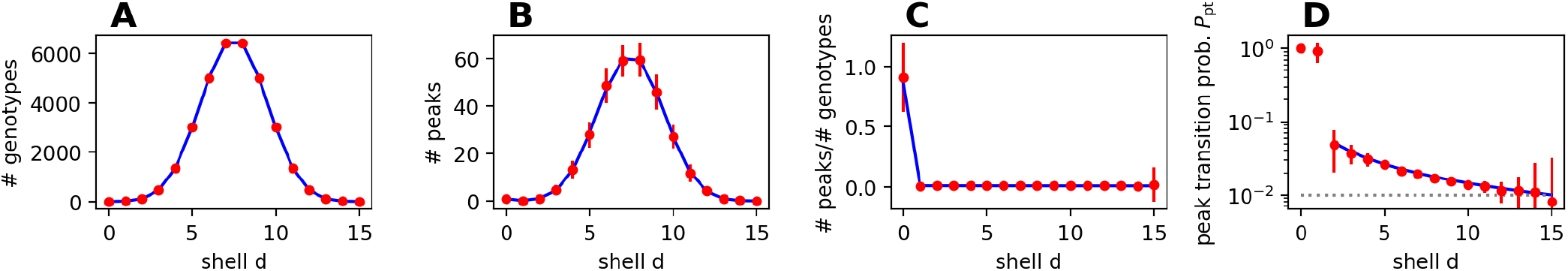
Number of genotypes, number of peaks, peak density and peak transition probability in each shell. *d*: In all plots, blue correspond to analytic results (eqs. 2-4) and red to simulation data (mean and standard deviation over 100 realisations of the sRMF model with *L* = 15 & *p* = 0.01). (A) Number of genotypes per shell as a function of distance to the reference genotype, *d*. (B) Number of peaks per shell. (C) Peak density, i.e. number of peaks normalised by the number of genotypes in the shell. (D) Probability of transitioning to a peak in the next step on a random adaptive walk. One mean *P*_pt_ value is computed for each shell in each landscape, and then the mean and standard deviation over the 100 landscapes is reported (landscapes with no non-peaks in shell *d* excluded). The analytic approximation from eq. 4 is not shown for shells *d≤* 1, which would require a careful treatment of the global peak position, and uses *n*_*d*_ = *L −d* since the mean is over all non-peak genotypes, not just those found on adaptive walks. The spike density *p* is shown as a baseline expectation (grey dotted line).

Next, we turn to peaks. Since peaks must be higher in fitness than all their mutational neighbours, only the reference genotype at *d* = 0 or a spike can be a peak. A spike genotype in shell *d >* 0 is a peak if it is a spike (with probability *p*) and all its *d* neighbours in shell *d* −1 are non-spikes (with probability (1 −*p*) ^*d*^). Similarly, the reference genotype in shell *d* = 0 is a peak if it is a spike (with probability *p*) or if neither itself nor any of its mutational neighbours are spikes (with probability (1 −*p*) ^*L*+1^). Thus, the expected number of peaks in shell *d* is:

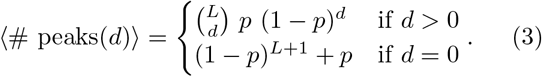

Like the number of genotypes, the number of peaks varies considerably from shell to shell, with a maximum in intermediate shells, see Fig 2B. This parallel between the number of genotypes and peaks per shell is not coincidental: Without the (1 −*p*)^*d*^-term, eqs. 2 & 3 would only differ by a constant factor, and one in 1*/p* genotypes in any shell with *d >* 0 would be a spike and a peak. Since (1 −*p*) ^*d*^ ≈1 in the small-*p* limit, this intuition remains useful, but is an overestimate, especially at high *d*.

Since the number of genotypes and peaks both vary from shell to shell, the shell with the highest *number* of peaks does not have to be the shell with the highest peak *density* relative to its number of genotypes, see Fig 2C.

### C. The shells with the highest number of peaks do not have the highest peak transition probability

Having described the location and density of peaks, we now turn to the probability that a random adaptive walk transitions to a peak in its next step, given that it is currently at a non-peak genotype in shell *d*. This genotype’s mutational neighbourhood can be characterized in two parts (assuming the low-*p* limit where every spike is a peak to good approximation):

- *L* −*d* neighbours in shell *d* + 1 (away from the reference genotype) [9]. Such a neighbour offers a fitness-increasing step to a peak if it is a spike, and a fitness-decreasing step otherwise. The probability that *j* of these neighbours are spikes is *B*(*j* |*n*_*d*_, *p*), where *B* denotes the binomial probability mass function and *n*_*d*_ = *L* − *d*. However, in the middle of a walk, the walker must have arrived at its current location via a fitness-increasing step, giving *n*_*d*_ = *L* − *d* − 1 unknown neighbours.
- *d* neighbours in shell *d* −1 (towards the reference genotype) [9]. Such a neighbour always has higher fitness. It is a peak if it is a spike. The probability that *i* of these neighbours are spikes is *B*(*i*|*d, p*).

Now, the probability of transitioning to a peak *P*_pt_ is the fraction of peaks among fitness-increasing neighbours, which we estimate as:

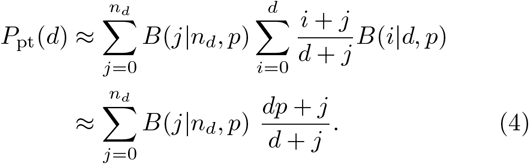

The simplification on the second line relies on the mean and sum of the binomial pmf.

Eq. 4 agrees with simulation data (Fig. 2D) and illustrates that *P*_pt_ increases towards the reference genotype: For large *d, n*_*d*_ is small and the sum is dominated by the *j* = 0 term, giving *P*_pt_ ≈*p*. For small *d*, non-spike uphill steps become rarer and *P*_pt_ *> p*. Thus, the peak transition probability, like the peak density, follows a different trend from the number of peaks per shell, see Fig. 2B&D.

The same arguments also define another probability that will be needed later: the probability of transitioning to a peak in shell *d* + 1 from a non-peak in shell *d*. By limiting the numerator to steps to shell *d* + 1, we approximate the probability for such a “step back”, *P*_sb_:

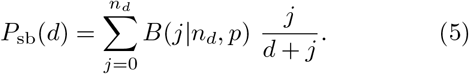

### D. Combining peak numbers, transition probabilities and path lengths gives the high-navigability phenomenon

The previous sections have illustrated that peak numbers are highest in intermediate-fitness shells, but that peak densities and peak transition probabilities follow a different pattern, decreasing with distance from the central shell. Building on this context, we now ask, how the highest-*x*% of peaks are reached on ≫*x*% of the walks. To address this, let us divide the landscape into a low-to-intermediate fitness region containing shells *d* ≥*d*_*c*_, which includes the high-peak-number shells and thus 100 −*x*% of all peaks. ≫*x*% of adaptive walks do not get stuck on these peaks and instead reach the *x*% of peaks in the high-fitness region. We hypothesise that this can be explained with the following two factors: First, unlike peak numbers, the probability of transitioning to a peak decreases with distance from the high-fitness region. Secondly, the low-to-intermediate fitness region is crossed by adaptive walkers in a small number of steps

since—if they do not transition to a spike—they only visit each shell once, even shells containing a huge number of genotypes. If the number of steps is small and the probability of transitioning to a peak at each step sufficiently low, then the probability of reaching the top-*x*% of peaks in the high-fitness region can be ≫*x*%. Note that both the *high* peak number and the *low* peak transition probability in the low-to-intermediate fitness region are important in the argument: if the peak number was lower, the fitness percentile of the high-fitness peaks would drop, and if the peak transition probability was higher, the probability of reaching the high-fitness peaks would drop.

To show that these features are sufficient to explain the high navigability of the sRMF model, we will turn this qualitative argument into analytic approximations. Thus, we will split the landscape at *d*_*c*_ and find expressions for two quantities: first, the share of high-fitness peaks and, secondly, the probability of ending at these high-fitness peaks. To approximate the share of high-fitness peaks, *x*, we use the expected number of peaks per shell (eq. 3):

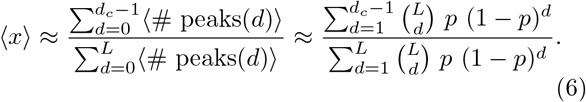

Next, we need *P* (end hf |*d*_*init*_), the probability of reaching these high-fitness peaks when starting from a non-peak genotype in shell *d*_*init*_. To compute *P* (end hf |*d*_*init*_), we would need to sum over two types of walks: walks stepping onto a spike *exactly* at *d* = *d*_*c*_− 1, and walks reaching non-spike genotypes at *d* = *d*_*c*_ − 1, which go onto peaks at *d* ≤ *d*_*c*_ − 1 if they do not take a step backwards to a spike in shell *d*_*c*_. For simplicity, we omit the first and likely smaller summand, thus obtaining a lower bound on *P* (end hf |*d*_*init*_). The second summand is given by the product of two probabilities: the probability of reaching shell *d* = *d*_*c*_− 1 without ending at a peak and the probability of not taking a step away from the reference genotype in the final step, to a peak in shell *d* = *d*_*c*_:

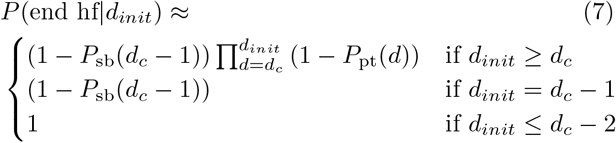

Here, *P*_pt_ is the probability of transitioning to a peak, given by eq. 4, and *P*_sb_ the probability of taking a step away from the reference genotype, given by eq. 5. We take *n*_*d*_ = *L*−*d*− 1 in both eq. 4 & 5, which is appropriate for adaptive walkers except in their first step.

Now, to get *P* (end hf), the probability of reaching high-fitness shells for an initial genotype sampled at random, we need to sum eq. 7 over initial shells *d*_*init*_, setting the probability of starting in shell *d*_*init*_ to the normalised number of genotypes in that shell (see eq. 2). Thus, neglecting the small probability that the initial genotype is itself a spike, we have:

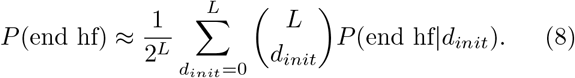

With eq. 8, we can approximate the probability of reaching the top *x*% of peaks, where *x*% depends on *d*_*c*_ through eq. 6. The resulting estimates for different values of *d*_*c*_ illustrate that the top *x*% of peaks are reached with probabilities ≫*x*%, as in the simulation data (blue line in Fig. 3). This is exactly the high-navigability phenomenon we were seeking to explain.

**Figure 3.**
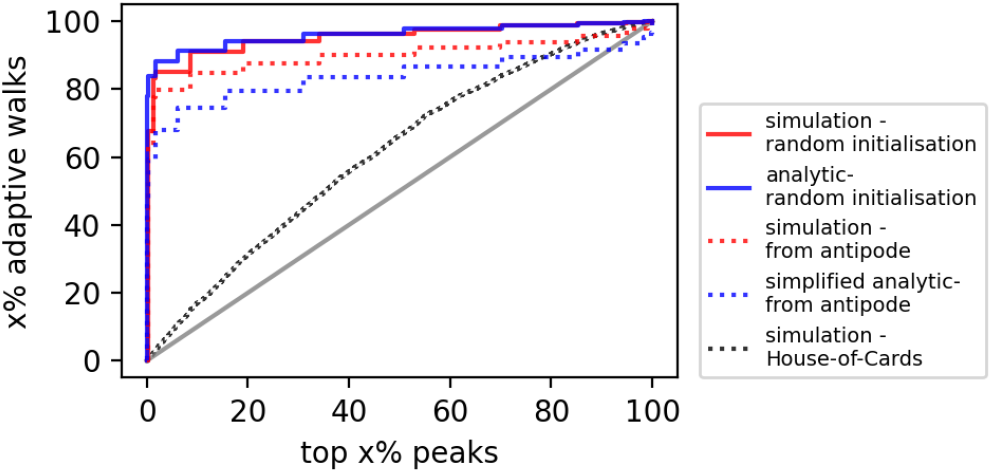
Analytic approximation of the sRMF landscape’s high navigability: Simulation results are shown in red for two scenarios: adaptive walks starting at a randomly chosen genotype in the landscape (solid line) and at the *d* = *L* shell, i.e. the antipode (dotted line). The corresponding analytic approximations are shown in blue: for each 2 ≤ *d*_*c*_ ≤ *L* − 1, the top-*x* % of peaks are computed from eq. 6 and plotted against the corresponding fraction of walkers approximated with eq. 8 (random initialisation)/eq. 9 (antipode). A grey line illustrates the naïve expectation, where *x*% of adaptive walks reach the top-*x*% of peaks, and a dotted black line shows simulations on an unstructured House-of-Cards landscape, where fitness values are independently drawn from a uniform distribution between 0 and 1. Both reference curves are exceeded by even the simplified analytic approximation from the antipode, implying that the landscape’s high navigability follows from the features accounted for in this approximation: low-to-intermediate fitness region with high # peaks, low *P*_pt_ and short paths. Simulations are based on a single landscape realisation (*L* = 15, *p* = 0.01) and 10^4^ adaptive walks.

To better relate the analytic approximation in eqs. 6–8 to our qualitative explanation at the start of the section, we will approximate a simplified lower bound for eq. 8, with the following assumptions: First, all walkers start in the shell furthest from the reference genotype, the antipode at *d* = *L*. Secondly, *P*_pt_ takes a constant value of *P*_pt_(*d*_*c*_) over all shells, a conservative estimate, which is higher than the true *P*_pt_(*d*) for *d* ≥ *d*_*c*_. Then we have:

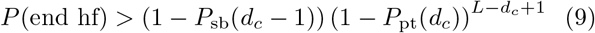

Here, the total number of factors, *L* −*d*_*c*_ + 1, is the number of steps from the antipode to the high-fitness region, (1 −*P*_sb_(*d*_*c*_ −1)) is the probability of not transitioning back to the low-to-intermediate fitness region, and *P*_pt_(*d*_*c*_) is the maximum peak transition probability in the low-to-intermediate fitness shells. Eq. 9 underestimates *P* (end hf) as expected, but still captures the high-navigability phenomenon (dotted blue line in Fig. 3): the share of adaptive walkers reaching the highest *x*% of peaks is ≫*x*% and higher than on an unstructured House-of-Cards landscape, where fitness is assigned randomly to genotypes and adaptive walks can reach slightly-higher-than-average peaks [18] (black dotted line in Fig. 3).

The simplicity of our final approximation in eq. 9 highlights key features which allow *P* (end hf) to be ≫ *x*%:

- First, the share of high-fitness peaks among all peaks, ⟨*x*⟩, is low because the shells with the highest number of peaks are of intermediate fitness.
- Secondly, the number of factors in eq. 9 is low because paths through the low-to-intermediate fitness region are short, of length *L* − *d*_*c*_ + 1.
- Thirdly, individual factors are close to one because the peak transition probability per step is low in the low-to-intermediate fitness region, i.e. *P*_pt_ ≪ 1, despite the high number of peaks in this region.

Thus, the analytic approximations are consistent with the qualitative explanation from the start of the section.

### E. Key features are also found in a highly navigable empirical landscape

Having analysed the high-navigability phenomenon in the sRMF landscape, we now turn to two versions of Papkou et al.’s [3] empirical *folA* landscape, which are both highly navigable [3, 7]: first, the full landscape and secondly, a sublandscape containing functional variants and their mutational neighbours (see Methods). Both landscapes are too complex to be approximated by simple mathematical expressions, but we *can* test on a qualitative level whether the central features continue to be present: a high number of peaks in the low-to-intermediate-fitness region yet a low peak transition probability, and a small number of steps for adaptive walkers to cross this region.

To investigate these three features, we first need to define a low-to-intermediate-fitness region in the *folA* landscape. For this, we follow the original paper [3] and denote peaks and genotypes as “high-fitness” if the second of the three mutated codons specifies an Asp or Glu amino acid. With this definition, both versions of the *folA* landscape have most of their peaks in the low-to-intermediate-fitness region (Fig. 4, first column), and this division perfectly separates a small number of high-fitness peaks from a large number of lower-fitness peaks [3]: high-fitness peaks are top-*x*% peak, with *x* = 1.8% in the full landscape and *x* = 14% in the sublandscape.

**Figure 4.**
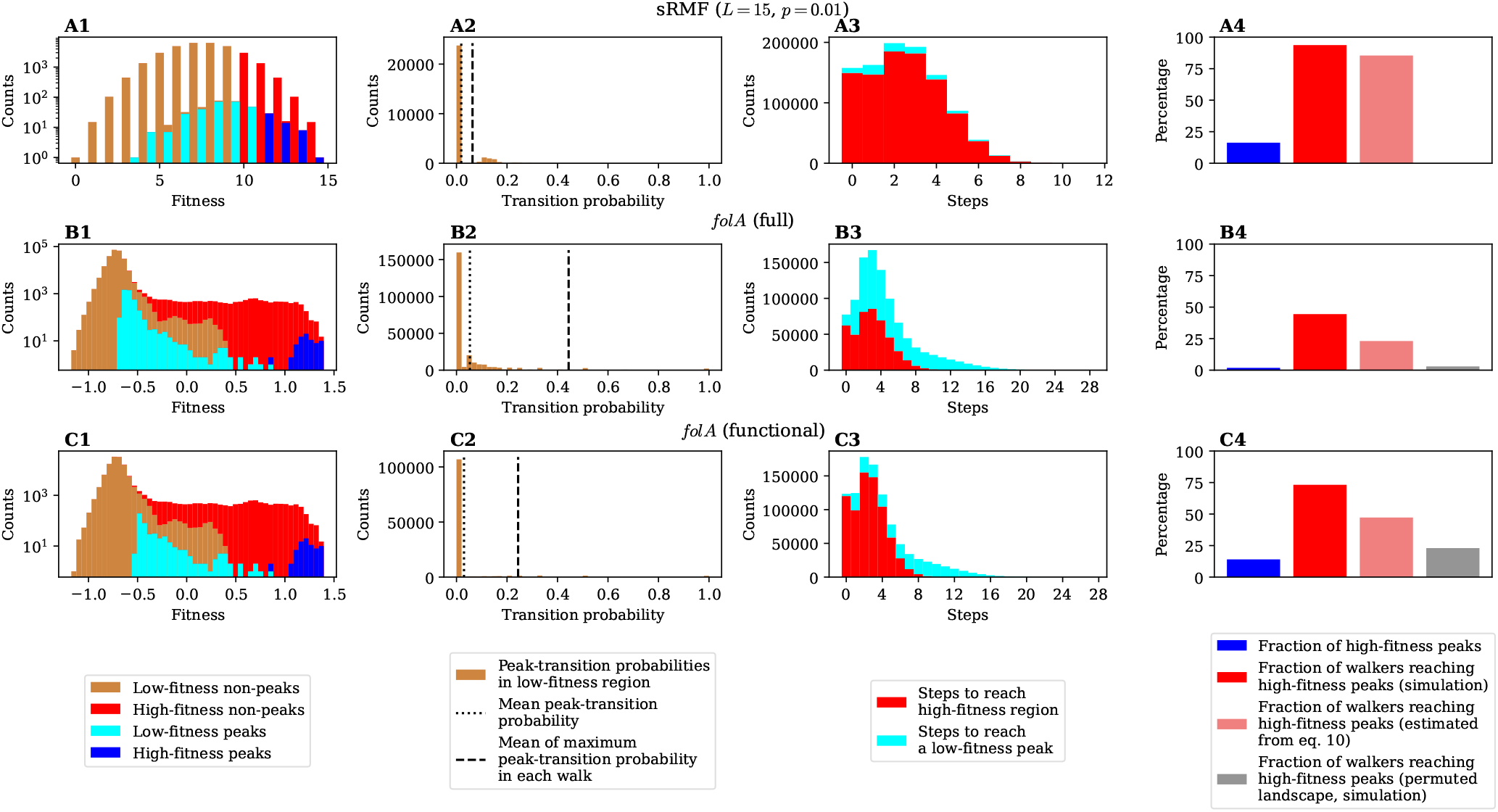
The low-to-intermediate-fitness region in the *folA* landscape shares the three features that we found to explain the high navigability in the sRMF landscape: The three rows contain data for: **(A)** The sRMF landscape. **(B)** The full *folA* landscape. **(C)** The functional *folA* sublandscape. **First column:** Stacked histograms for non-peak and peak genotype fitness, showing that most peaks are in the low-to-intermediate-fitness region. **Second column:** The probability of transitioning to a peak, *P*_pt_, is shown as a histogram for all non-peak genotypes in the low-to-intermediate-fitness region. The dotted line represents the mean; the dashed line first takes the maximum *P*_pt_ over the low-to-intermediate-fitness-region segment of each adaptive walk and then the mean over walks. **Third column:** Stacked histograms of the number of adaptive walk steps until reaching either the high-fitness region or a low-to-intermediate-fitness peak, whichever occurs first (zero steps implies the initial genotype is a peak or in the high-fitness region). **Fourth column:** Bar plot quantifying navigability. The bars show the fraction of peaks found in the high-fitness region (blue; this is equivalent to their fitness percentile from the top, *x*%), the fraction of simulated adaptive walks reaching these high-fitness peaks (red), the estimated fraction of adaptive walks reaching these high-fitness peaks (pale red, eq.10), and the fraction of walkers reaching the same fraction of highest-fitness peaks in simulations on a permuted version of the landscape (grey). We find that the approximation from eq.10, which is only based on *P*_pt_ and path lengths, already predicts high navigability, with≫ *x*% of walks reaching the top-*x*% of peaks. The plots are based on 10^6^ simulated adaptive walks. We omit the permuted sRMF landscape because the neutral mutations resulting from discrete fitness values would require a special treatment.

For non-peak genotypes, the division is no longer perfect and there is a fitness range, for which genotypes can lie in both the low-to-intermediate-fitness and the high-fitness region (Fig. 4, first column). Despite this overlap, adaptive walks almost never revert from the high-fitness region to the low-to-intermediate-fitness region (probability *<* 0.1% in both landscapes). Thus, just as in the sRMF model, adaptive walkers only need to have a ≫*x*% probability of leaving the low-to-intermediate-fitness region without being trapped at a peak, and we have the high-navigability phenomenon. This probability, *P*_end hf_, is indeed ≫*x*% in both landscapes (see Fig. 4, blue and red bars in fourth column). The question is whether this high *P*_end hf_ could be explained by similar landscape features as in the low-to-intermediate-fitness region of the sRMF model: low peak transition probabilities and short path lengths crossing the region. The second and third columns of Fig. 4 show that, on a qualitative level, these features are present in the *folA* landscapes, consistent with the short path lengths found in the original paper [3]. One notable difference with the sRMF model lies in a longer-tailed path length distribution of paths ending at low-to-intermediate-fitness peaks, which is due to one special trapping region among the low-to-intermediate-fitness genotypes (see SI).

Going beyond the qualitative similarity between the sRMF and *folA* landscape features, we tested whether the order of magnitude of the path length and *P*_pt_ val-ues is sufficient to imply a high probability of reaching the high-fitness region. To do so, we selected a representative peak transition probability *P*_pt_ by averaging over the maximum peak-transition probability in the low-to-intermediate-fitness segment of each adaptive walk. This is a more conservative choice than the mean peak transition probability *P*_pt_ (see Fig. 4, second column) and was selected to account for high-*P*_pt_ outliers. Then, we approximate the probability of ending at the high-fitness peaks in analogy with the sRMF model as:

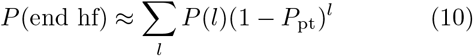

Here, *P* (*l*) is the share of walks with *l* steps, either to the high-fitness region or a low-to-intermediate-fitness peak. While the estimate from eq. 10 is lower than the simulated probability of reaching high-fitness peaks, it is much higher than the share of high-fitness peaks and thus captures the high-navigability phenomenon (pale red bar in the fourth column of Fig. 4). This estimate from eq. 10 is also higher than the baseline from a permuted ‘House-of-Cards’ [1] fitness landscape, where the genotype-fitness assignments in the landscape are shuffled (grey bar in Fig. 4, fourth column). Thus, while the estimate is far from exact, the ≫*x*% share of walks ending at high-fitness peaks can be explained based on just three numbers: first the low peak transition probabilities in the low-to-intermediate-fitness region, secondly the high share of peaks in the low-to-intermediate-fitness region and thirdly the short walk lengths to the high-fitness region.

To sum up, the features of the empirical *folA* landscape are consistent with the hypothesis that the high-navigability phenomenon in this landscape can be explained with similar arguments as in the sRMF model. However, unlike the sRMF model, the insights derived from analytic approximations have to be taken with caution, since they average over a highly heterogeneous region in the landscape.

### F. Relationship between peak fitness and basin sizes

Finally, let us take an alternative perspective on the end point of adaptive walks, by following the original *folA* paper [3] and focussing on basins: A genotype *g* is in a peak *s*’s basin if a fitness-increasing accessible path exists from *g* to *s*. A genotype can belong to the basins of multiple peaks, so being in *s*’s basin only means that *s can* be reached, without giving the probability that *s will* be reached. Thus, we need two quantities for each peak: its normalised basin size, i.e. the fraction of all genotypes falling within the basin of that peak, and the probability that the peak will be reached by an adaptive walker starting at an arbitrary genotype in its basin. Then, if an adaptive walk starts at a randomly chosen genotype, the probability that it ends on a given peak *s* can be written as:

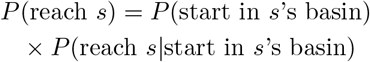

*P* (reach *s*) is orders of magnitude lower for many low-to-intermediate-fitness peaks than for the highest-fitness peaks in both the sRMF and *folA* landscapes (Fig. 5), consistent with the high navigability of these landscapes. When decomposing *P* (reach s) into *P* (reach s start in s’s basin) and *P* (start in s’s basin), we find that both factors play a role in suppressing low-fitness peaks: peaks with a low probability of being reached can either have small basins or a low probability of being reached from their basin, or both. Normalised basin sizes *P* (start in s’s basin) tend to increase with peak fitness, as previously noted [3, 7]^2^. However, *P* (reach s|start in s’s basin) also varies by orders of magnitude from peak to peak, and thus knowing two basin’s sizes is insufficient for computing their relative likelihood of being reached. This is consistent with Li and Zhang’s [7] finding that basin sizes alone, specifically their correlation with peak fitness, are not the most relevant quantity for predicting the percentage of walkers ending at the top-14% of peaks.

**Figure 5.**
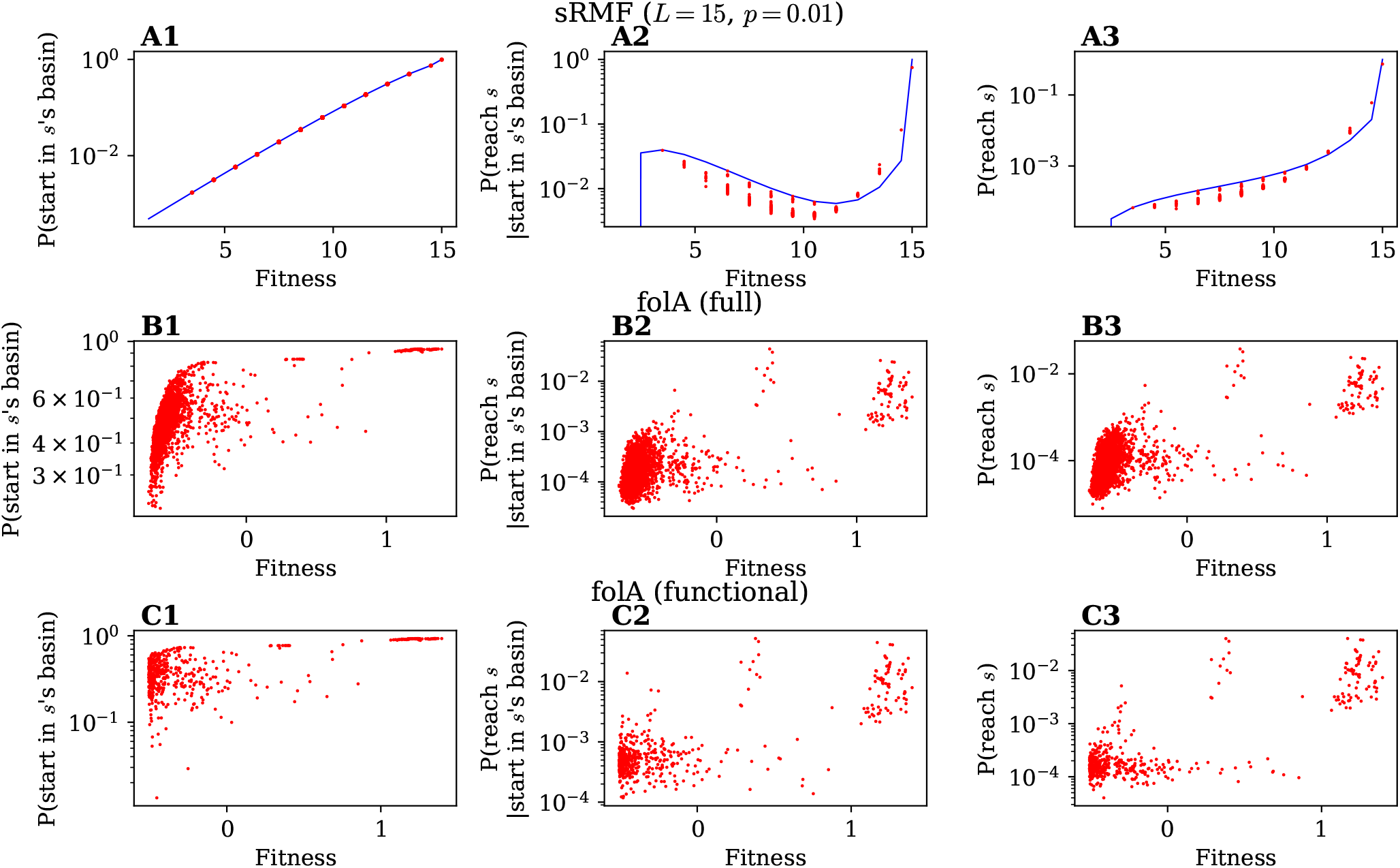
Dependence of basin properties on peak fitness: The three rows contain data for: **(A)** The sRMF landscape. **(B)**The full *folA* landscape. **(C)** The functional *folA* sublandscape. **First column:** basin size (normalised by the total number of genotypes in the landscape) against peak fitness. **Second column:** *P* (reach s |start in s’s basin) against peak fitness. **Third column:** *P* (reach s) against peak fitness. Again, simulations are shown in red, analytic approximations for the sRMF model in blue (eqs. 11 and eq. 12). sRMF simulations are based on a single landscape realisation (*L* = 15, *p* = 0.01). While basin sizes are exact, the probabilities in the second and third columns are estimated from 10^6^ adaptive walks initialised on randomly chosen genotypes in the landscape.

To gain insight into the peak-height-dependence of both factors in the sRMF model, we focus on the low-*p* limit, where spikes are rare. Thus, we approximate the sRMF landscapes by an almost smooth Mount Fuji, where the focal peak *s* is the only spike in an otherwise purely additive landscape. In this landscape, an accessible path exists from *g* to *s* if all steps from *g* to *s* mutate a ‘1’ to a ‘0’, as on a perfectly smooth Mount Fuji [12], with one exception: a single mutation from ‘0’ to ‘1’ is possible in the final step due to the elevated height of the focal spike *s*. Thus, if our single spike *s* is in shell *d*_*s*_, genotypes in the basin of *s* have to fulfil the following criteria: they have complete freedom in the *L* −*d*_*s*_ positions that are ‘0’ in *s*, but the remaining *d*_*s*_ positions all have to be ‘1’ with at most a single exception. These conditions are satisfied by 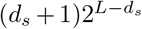 genotypes. Normalising by the number of genotypes in the landscape, 2^*L*^, we have the following normalised basin size, *P* (start in s’s basin):

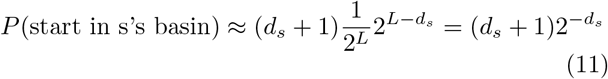

Thus, basin size increases exponentially with decreasing *d*_*s*_ and thus with increasing fitness.

Continuing with the approximation, where the focal peak *s* is the only spike in an otherwise additive landscape, we now turn to the probability of reaching *s* in an adaptive walk starting from a genotype *g* in the basin of *s*. The combinatorics of adaptive walks from *g* to *s* depends on whether a 0 →1 mutation is needed at the last step. Here, we will assume that one such mutation is needed since this applies to *d*_*s*_*/*(*d*_*s*_ + 1) of the genotypes in the basin, which is more than half the genotypes if *d*_*s*_ *>* 1. In this case, adaptive walks from *g* reach *s* if:

1. Out of all the *d*_*g*_ uphill mutations from *g*, those position in which *g* has a 1 and *s* has a 0 mutate first. Since the 1’s in *g* can mutate to 0’s in any order, one in 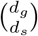 paths fulfils this criterion.
2. The correct 0 → 1 mutation happens at the last step. Since the spike is one of *d*_*s*_ fitness-increasing neighbours at the final step, this probability is 1*/d*_*s*_

Thus, for *d*_*s*_ *>* 1, we have:

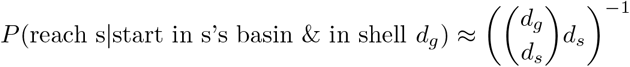

To remove the dependence on the initial shell *d*_*g*_, we need to sum over *d*_*g*_, weighted by the number of genotypes in the basin for each *d*_*g*_. Let us focus on all genotypes with the 0→ 1 mutation at the same fixed position. In total, there are 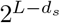 such genotypes in the basin, of which 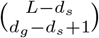 are at distance *d*_*g*_ from the reference genotype since these genotypes must have *d*_*g*_ −*d*_*s*_ + 1 0 →1 flips compared to the peak *s* together with their 1→ 0 flip. Since we are assuming that the initial genotype has one ‘0’ at a position where *s* has a ‘1’, these genotypes lie between shells *d*_*s*_ and *L* − 1, giving:

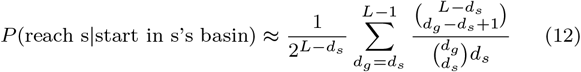

The expressions derived so far only apply to spikes with *d*_*s*_ *>* 1. For the innermost shells with *d*_*s*_ ≤ 1, we note that a landscape cannot have peaks in both shells *d* = 1 and *d* = 0. In our case of *L* = 15 & *p* = 0.01, a peak in shell *d* = 0 is more likely. In our single-spike approximation, this global peak has all genotypes in its basin and is reached by all accessible paths, thus giving *P* (reach s|start in s’s basin) ≈ *P* (start in s’s basin) ≈ 1.

Despite the approximations underlying eqs. 11&12, they approximate the simulation data well (Fig. 5, first row). In both the simulation data and the analytic approximation, we find low-fitness peaks to be suppressed by low normalised basin sizes, *P* (start in s’s basin), and intermediate-fitness peaks by a low probability of being reached from the respective basin, *P* (reach s |start in s’s basin). This suppression of intermediate-fitness peaks can again be understood in terms of peak densities in different shells: Intermediate-fitness peaks are found in large shells and these shells could be crossed via many alternative genotypes, depending on the order in which mutations are accumulated, causing the probability of taking the path via peak *s* to be low.

## III. CONCLUSION AND DISCUSSION

In this paper, analytic approximations and computer simulations on a simplified Rough Mount Fuji model have illustrated, how a fitness landscape can be highly navigable, such that the top-*x*% of all peaks are reached with a probability greatly exceeding *x*%. The sRMF landscape has the highest number of peaks as well as the highest number of genotypes at intermediate fitness, but normalised quantities like peak density and the probability of transitioning to a peak follow a different trend. Thus, the peak transition probability at low-to-intermediate fitness is low enough that adaptive walks are unlikely to be trapped at an intermediate-fitness peak, as long as they follow short paths to the highest-fitness genotypes, guided by the additive fitness gradient. Then, the fraction of adaptive walkers ending in the high-fitness region far exceeds the share of peaks found in this region. Note that both the *high* peak number and the *low* peak transition probability in the low-to-intermediate fitness region are important: if the peak number was lower, the fitness percentile of the high-fitness peaks would drop, and if the peak transition probability was higher, the probability of reaching the high-fitness peaks would drop. The fact that short paths *can* cross a large low-to-intermediate fitness region is due to the high dimensionality of fitness landscapes, which is easily neglected in our low-dimensional intuition [17]. While our mathematical treatment only applied to the sRMF model, we found similar features in the empirical *folA* landscape, suggesting that similar principles may apply. Finally, we revisited the likelihood of reaching high-fitness peaks from a basins perspective, relying on two quantities: basin sizes and the probability of reaching a peak from its basin.

The sRMF model mirrors several features of the standard RMF model found in previous calculations: Peak density decreases with distance to the reference genotype [9], the highest peak number is found at intermediate distance to the reference genotype [9], the high-fitness reference sequence is likely to have an accessible path from most genotypes [13, 14] and adaptive path lengths can reach order *L* ∼[15]. Both the standard RMF model and the sRMF model are idealised and describe landscapes with a linear fitness gradient towards a reference genotype. The sRMF is even more idealised, but its simplicity facilitates mathematical and conceptual analyses, making it a useful tool for addressing further questions about multi-peaked fitness landscapes.

Our analysis agrees with additional results in the larger field of fitness landscapes: First, our basin analysis highlights that the probability of staring in a peak’s basin is insufficient for approximating the probability of reaching that peak, reflecting fact that a peak is not guaranteed to be reached even when an accessible path exists [15] and highlighting the importance of going beyond accessibility analyses. Secondly, the central role of peak densities in our analysis mirrors existing research, for example on adaptive path lengths [19, 20].

Our calculations offer two explanations of a seemingly counter-intuitive result: how the fitness distribution of adaptive walk endpoints can differ so drastically from the fitness distribution of peaks in a fitness landscape. However, even within the simple sRMF model, our two perspectives—on peak densities and basin sizes—may only be two of many ways of breaking down the high-navigability phenomenon. Whether different principles are needed for other highly navigable landscapes is an open question.

One caveat concerns our treatment of the empirical *folA* landscape: We took the landscape at face value since our objective was to clarify previous work [3, 7] by showing how adaptive walkers reach the highest peaks in this landscape. However, when accounting for experimental noise, the true number of peaks may be lower [21, 22].

Another limitation is that we focus only on random adaptive walks, where all fitness-increasing mutational steps are equally likely. The opposite limiting case of greedy adaptive walks would give deterministic dynamics and could thus be addressed with a basin perspective. These basins would be computed based on deterministic greedy trajectories and thus differ from the basins in the present paper. Future work should analyse such greedy dynamics and relax the strong-selection-weak-mutation assumption in favour of more complex models of evolving populations, where outcomes can depend on population-genetic properties like population sizes [23, 24].

Finally, our analysis has focused on a relatively short sequence length of *L* = 15, giving a landscape of ≈3 ×10^4^ genotypes, similar to the ∼ 10^5^ genotypes in the *folA* landscape. Future work could use the mathematical tractability of the sRMF model to also investigate longer sequence lengths of *L* ≈ 50 – meaning 10^15^ genotypes, beyond the realm of exhaustive genotype-to-fitness mappings.

## IV. METHODS

### A. Peak percentiles

In the sRMF model, we commonly have several peaks of equal fitness. Thus, to compute the peak percentile corresponding to a given fitness, we take a conservative approach and define the percentile from the top as the fraction of peaks with higher *or equal* fitness. For example, if the peak heights in the landscape were (9, 8, 8, 8,

5, 5, 5, 5, 4, 3), peaks of height 8 would be among the top-40% of peaks and percentiles in the range 10–39% would only include the peak of height 9. For corresponding percentiles of adaptive walks, we compute the fraction of walks that reach a peak with percentile *x*% or better.

### B. Basins

Basins are defined, such that a genotype is in a peak’s basin if it has an accessible path to this peak [3, 7]. To identify all genotypes in a peak’s basin, we start with a sparse matrix containing the genotype-to-genotype transition probabilities for a random adaptive walk. Then, we start with the peak genotype and perform a recursive breadth-first search of non-zero incoming transitions, until no new genotypes are added to the basin.

### C. *folA* landscape

The data defining the *folA* landscape is read in from Papkou’s [25] supplementary data file `in_data/fitness_data_wt.rds`. For the full land-scape, we take all 261, 333 genotypes with fitness measurements, giving 4055 fitness peaks. For the sublandscape, we follow Papkou et al. [3] and reduce the landscape to the set of all functional genotypes with fitness ≥ −0.508 and their neighbours; we find that the resulting landscape consists of a single connected component, containing 135, 559 genotypes, and 527 fitness peaks, differing slightly from the 135, 178 and 514 in the original analysis by Papkou et al. [3].

### D. Permuted landscapes

Permuted landscapes are generated by shuffling the fitness values between all genotypes contained in a landscape. Then, we simulated adaptive walks on the permuted landscapes and recorded the fraction of walks ending at high-fitness peaks, i.e. the top-*x*% of peaks, where *x* is set to match the value in the corresponding unpermuted landscape. Since peak numbers are finite, a perfect match is not possible and *x* is set to be as close as possible to the target value without exceeding it.

## V. ACKNOWLEDGEMENTS

We acknowledge support of the Spanish Ministry of Science and Innovation through the Centro de Excelencia Severo Ochoa (CEX2020-001049-S, MCIN/AEI /10.13039/501100011033), the Generalitat de Catalunya through the CERCA programme and the EMBL partnership. Research for this publication has been carried out in the Barcelona Collaboratorium for Modelling and Predictive Biology. We thank M. Giraud for feedback.

## Footnotes

1 This convention follows the title of the *folA* paper [3], but differs from other uses of the word *navigable* as a synonym for *accessible* [5].

2 Our basin sizes in the sublandscape differ from Papkou et al. [3], who focused on functional genotypes only. We include all genotypes in the sublandscape for consistency with the adaptive walks, which can start at any genotype in the sublandscape.

