## Supplementary Information for "Using a simplified Rough Mount Fuji model to disentangle how multi-peaked fitness landscapes can be highly navigable"

### Contents

**S1 Additional tests using modified versions of the sRMF model** **1**

**S2 Additional data on adaptive walks on the empirical folA landscape** **3**

### S1 Additional tests using modified versions of the sRMF model

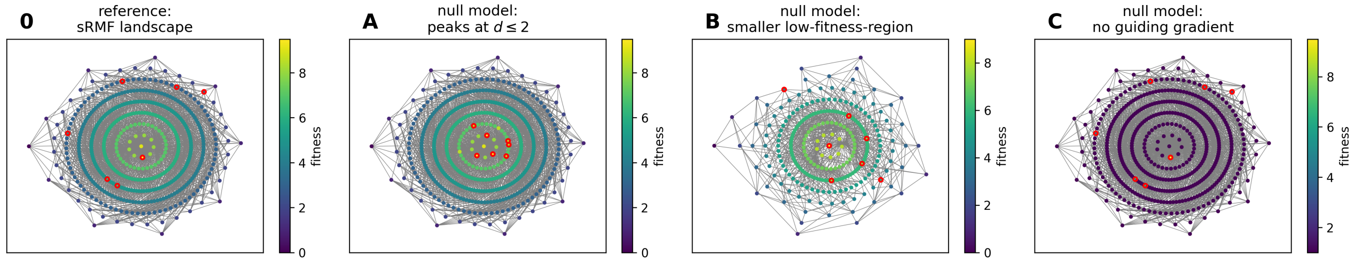


Figure S1: **Construction of null models perturbing the three features of the sRMF model that we identified to be relevant for its high navigability:** 0) sRMF landscape as a basis for the three null models. A) Null model with spikes only in high-fitness shells and same total peak number as corresponding sRMF landscape. B) Null model with smaller shells, especially in the low-to-intermediate-fitness region, and same total peak number as corresponding sRMF landscape. C) Null model with same peaks and same peak fitness as in corresponding sRMF landscape, but otherwise constant fitness and no guiding gradient towards the reference sequence. For the purpose of visualisation, the parameters are not the same as in the simulations in Fig. S2: $L = 9$ and $p = 0.02$, with $d_{\text{peaks}} = 2$ in null model A.

In the main text, we identified three features, which allow the sRMF landscape to be highly navigable: a high number of peaks in the low-to-intermediate-fitness region, yet a low peak transition probability, and a fitness gradient guiding populations towards the high-fitness region in few steps. Here, we create modified versions of the sRMF model to further analyse the role these three features play for the high navigability of the landscape (as illustrated in Fig. S1). Each null model is defined relative to a sRMF landscape, and matches the number of peaks in this sRMF landscape:

A) **Null model without high number of spikes in low-to-intermediate fitness region**: We start with a smooth Mount Fuji without any spikes and then add spikes for randomly drawn high-fitness sequences only ($d \leq d_{\text{peaks}}$ from the reference sequence), until reaching the target number of peaks.

B) **Null model with smaller shells, especially in the low-to-intermediate-fitness region**: We start with a smooth Mount Fuji without any spikes and reduce its number of genotypes, especially in the low-to-intermediate-fitness region: genotypes are only included in the landscape if their "0"s are within the first $D + 3$ positions, where $D$ is the distance from the antipode. We choose this structured reduction of genotypes over a random one to ensure that each genotype still has mutational neighbours in both its adjacent shells. Then we add spikes for randomly drawn sequences until the number of peaks matches that of the corresponding sRMF model. To match the number of peaks in the smaller landscape, we use a lower peak density of $p = 0.001$ in the corresponding sRMF model.

C) **Null model without a guiding gradient towards the reference sequences**: To create a null model, we take the corresponding sRMF landscape and set the fitness of all non-peak genotypes to $F_{\text{non-spike}} = 1$. This value ensures that non-peaks remain lower than the lowest peaks, with fitness 1.5, and thus leave the peak identities intact. Thus, the peaks and their fitness values match the sRMF landscape exactly, but the landscape is otherwise flat. We allow the random adaptive walks to take fitness-preserving neutral steps with the same probability as uphill steps.

The results of all three tests are shown in Fig. S2. While it is not possible to perfectly isolate the three features (for example, changes to the peaks affects both peak transition probabilities and the number of peaks in the low-to-intermediate-fitness region), the tests further support our conclusions from the main text: All three null models are less navigable than the corresponding sRMF landscapes, and the three features perturbed by the null models thus contribute to the high navigability of the sRMF model.

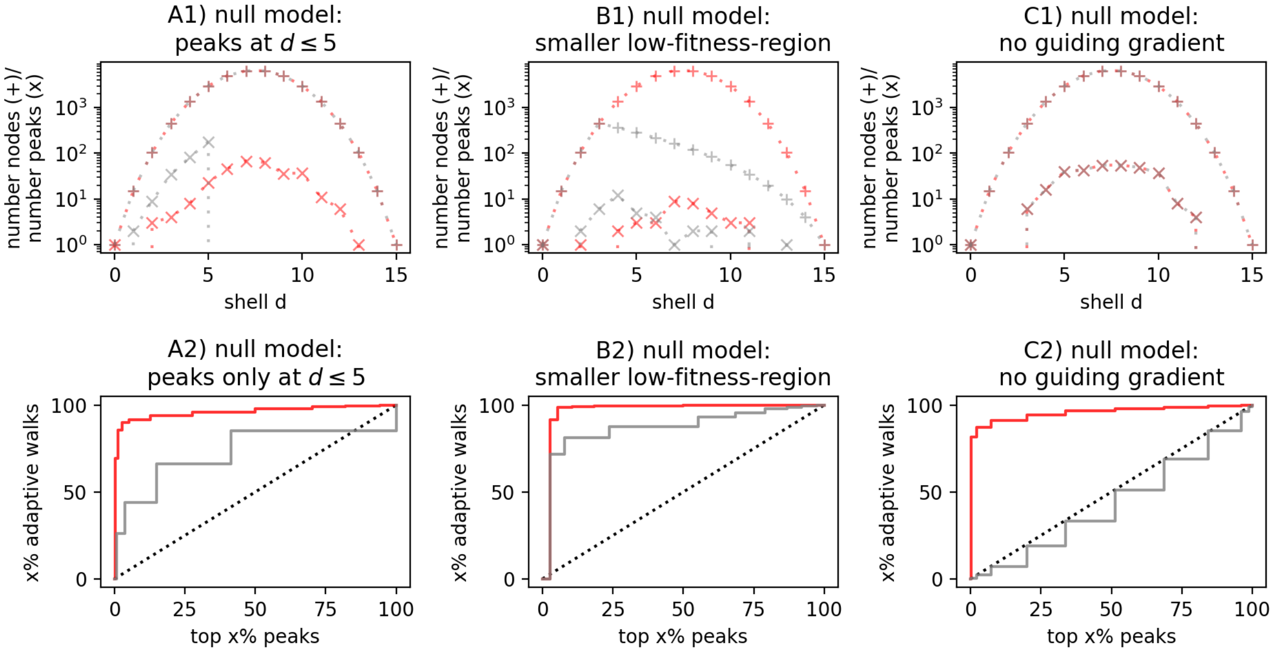


Figure S2: **Analysis of null models perturbing the three features of the sRMF model that we identified as relevant for its high navigability: Columns A - C:** Each column focuses on one of the three null models described in the text. For each null model, we first create one sRMF landscape and match the total number of peaks in the null model ($L = 15$, $p = 0.01$ for null models A&C, $p = 0.001$ for null model B). **Row 1**: number of genotypes ('+'-markers) and number of peaks ('x'-markers) per shell, for both the sRMF model (red) and the corresponding null model (grey). This shows that the null models are successfully constructed: the first null model has peaks only for $d \leq 5$, the second null model has fewer genotypes, especially in the low-to-intermediate-fitness region, and the final null model has the same number of genotypes and peaks per shell as the corresponding sRMF model. **Row 2**: The percentage of $10^4$ simulated random adaptive walks reaching the top-$x\%$ of peaks is plotted for each null model (grey) and its corresponding sRMF model (red). The null models have lower navigability, implying that the features they perturb are indeed important for navigability.

### S2 Additional data on adaptive walks on the empirical folA landscape

In the main text, we approximated the share of adaptive walks reaching high-fitness peaks in the empirical *folA* landscape from refs [1, 2] as $P(\text{end hf}) \approx \sum_l P(l)(1 - P_{\text{pt}})^l$. This expression assumes that averages are sufficient to capture the heterogeneous low-to-intermediate-fitness region of the landscape. To test the validity of this assumption more carefully, we perform the following two analyses (Fig. S3):

1. To test whether adaptive walks of different lengths experience similar peak transition probabilities, we stratify the peak transition probabilities encountered on simulated adaptive walks by walk length. We find no strong trend among the path lengths $l$ with high $P(l)$, indicating that walk length and peak transition probability can be assumed to be independent to first approximation.

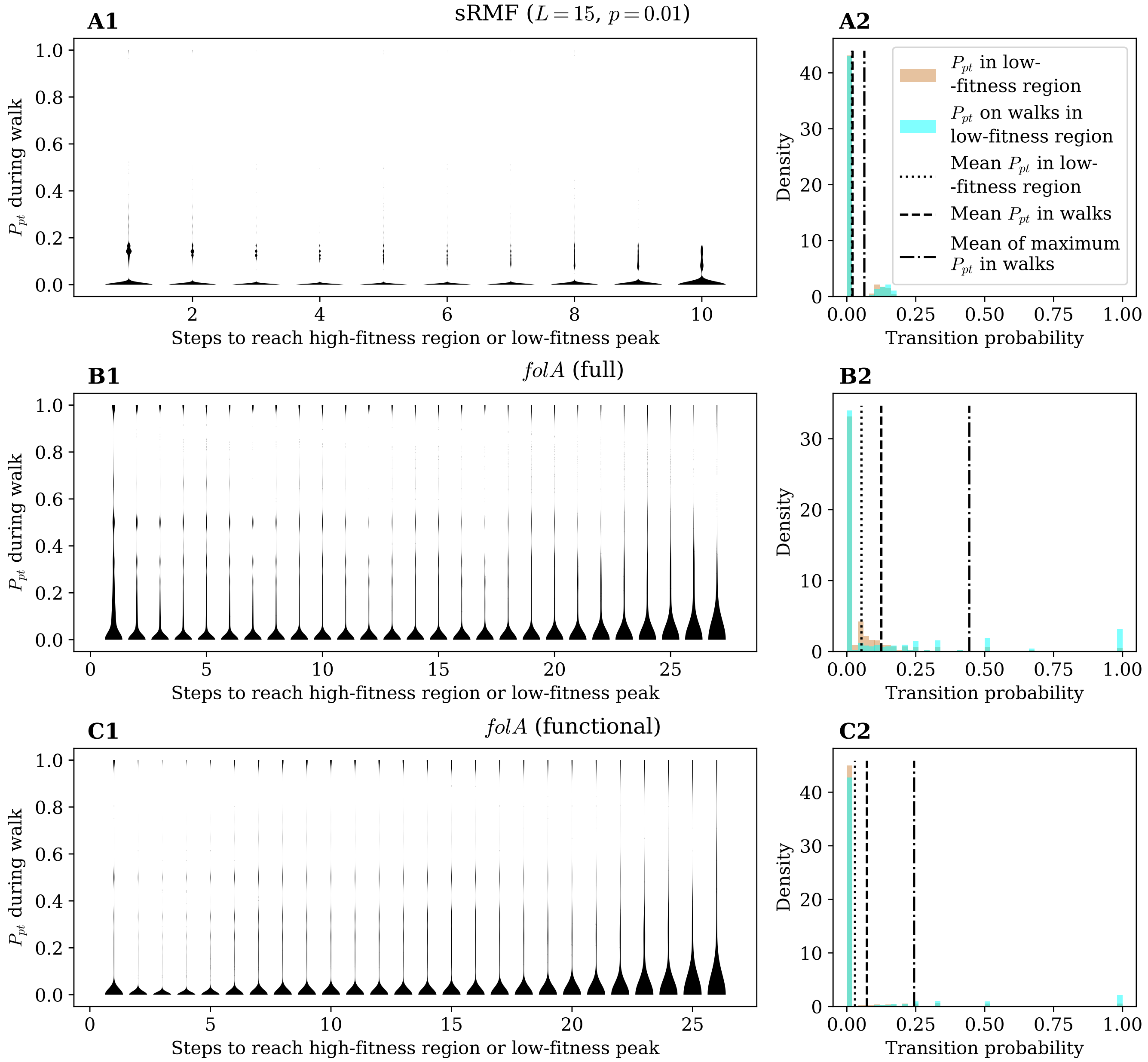


Figure S3: **Additional adaptive walk data for the landscapes studied in this work. Left column:** The distribution of the peak-transition probabilities seen by walkers until entering the high-fitness region or being trapped at a peak, stratified by the number of steps taken to reach the high-fitness region or a peak. **Right column:** The distributions of peak-transition probabilities taken across all non-peak genotypes in the low-to-intermediate-fitness region, and across genotypes collected from the low-to-intermediate-fitness segments of $10^6$ adaptive walks (excluding peak genotypes). Included are lines showing the mean values of both distributions, and the mean of the maximum peak-transition probability seen in each walk before entering the high-fitness region or being trapped at a peak (i.e. the value of $P_{pt}$ used in eq. 10). In plot **A2**, the dotted and dashed lines coincide almost exactly.

2. To test whether adaptive walks lead through genotypes with above-average or below-average peak transition probabilities, we compare the distribution of peak transition probabilities over all genotypes in the relevant low-to-intermediate-fitness region to that over the low-to-intermediate-fitness genotypes found along adaptive walks. Our qualitative conclusion – that peak transition probabilities in the low-to-intermediate-fitness region are low – remains unaffected.

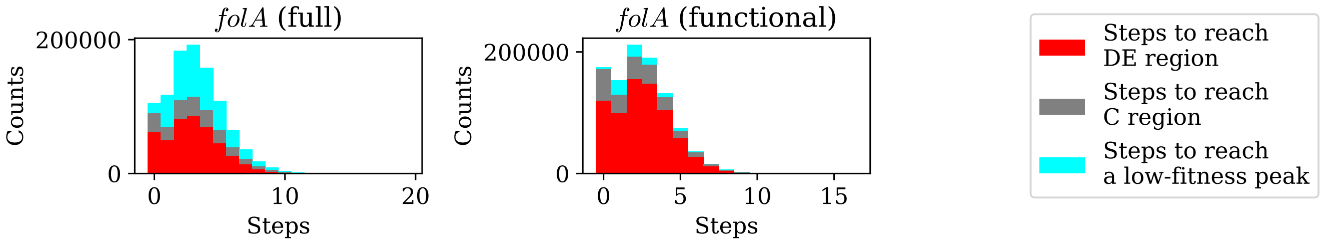


Figure S4: **Long walks in *folA*'s low-to-intermediate-fitness region quickly fix cysteine.** Compared with the corresponding histograms in Fig. 4 in the main text, the number of steps in adaptive walks is shown here until the walks enter the high-fitness region (i.e. fix aspartate or glutamate in position 27, in red), *or* fix cysteine in position 27 (grey), or are otherwise trapped at a peak (cyan).

Moreover, we turn to one special feature of the path length distribution in the *folA* landscapes: the number of steps to reach a high-fitness peak tends to be lower than the number of steps to reach a peak in the low-to-intermediate-fitness region. This can be explained as follows: In the *folA* landscapes, typical adaptive walks quickly step onto a genotype coding for an Asp/Glu or Cysteine amino acid at the second codon. If the genotype codes for Asp/Glu, the high-fitness region is reached, and the walk length is short. If, however, the genotype codes for Cysteine, then the walk enters an isolated part of the low-to-intermediate-fitness region, from which the high-fitness region is unlikely to be reached, but where adaptive walks can continue for some time until ending at a low-to-intermediate-fitness Cysteine peak. Even though the Cysteine peaks only constitute a small fraction of low-to-intermediate-fitness peaks (1.26% in the full landscape, 8.16% in the sublandscape), the other low-to-intermediate-fitness peaks are so unlikely to be reached that paths to the Cysteine peaks dominate the walk length statistics. Thus, the long path lengths in the Cysteine region dominate the statistics. This interpretation is supported by the following analyses (Fig. S4):

1. Once a Cysteine genotype fixes, only a small fraction of adaptive walks reach the high-fitness region (0.23% in the full landscape, 0.22% in the sublandscape). Thus, the Cysteine genotypes, while part of the low-to-intermediate-fitness region, form a trapping region, from which progression to the high-fitness region is unlikely.

2. Once we change the endpoint of the walks - either to the first low-to-intermediate-fitness peak, *or to a Cysteine genotype* or to the high-fitness region - the path length distribution for walks ending at low-to-intermediate-fitness peaks loses its pronounced long-$l$ tail. Thus, one of these three milestones occurs quickly, and the long tail was due to trapping in the Cysteine region.
